# Estimating Biochemical Concentration in Food Using Untargeted Metabolomics

**DOI:** 10.1101/2022.12.02.518912

**Authors:** Michael Sebek, Giulia Menichetti, Albert-László Barabási

**Author notes:** These authors contributed equally to this work (Michael Sebek, Giulia Menichetti).

## Abstract

Untargeted metabolomics can detect hundreds of biochemicals in food, yet without standards, it cannot quantify them. Here we show that we can take advantage of the universal scaling of nutrient concentrations to estimate the concentration of all biochemicals detected by untargeted metabolomics. We validate our method on 20 raw foods, finding an excellent agreement between the predicted and the experimentally observed concentrations.

---

The documented impact of diet on health has prompted national efforts to catalogue the biochemical composition of food^1^, like the USDA Standard Reference^2^ that reports the concentration of 150 nutrients in 9,759 foods. The data was collected using gravimetry (AOAC934.06), liquid chromatography (AOAC988.15), and fluorimetry (AOAC967.22), experimental methodologies that can detect and quantify the concentration of a biochemical in food. Meanwhile untargeted metabolomics can detect hundreds more compounds within a single experiment, but only relative concentration are inferred, mainly because of the unknown fraction of molecules ionized for each compound^3,4^. Targeted metabolomics circumvents ionization efficiency^5^ by using standard curves to determine the concentration, a low-throughput procedure that eliminates the advantages of untargeted metabolomics over established methodologies by increasing the time and cost over current AOAC methods. Untargeted metabolomics needs to move beyond relative concentrations for adoption by food composition catalogues. Here we report a method to predict concentrations from untargeted metabolomics, offering researchers a way to extend their results beyond relative concentrations. This is advantageous for food composition catalogues which can leverage the predictions to report expected concentrations and their potential variance for foods in a high-throughput way.

We show that the universal scaling of nutrient concentrations across the food supply ^6^ allows us to determine nutrient concentration in untargeted metabolomics. We performed metabolomics experiments on 20 raw plant ingredients to minimize the influence of human processing and span the phylogenetic tree as well as edible parts of plants (fruits, leaves, roots) to get a representative diversity of plant composition across the food supply (SI Experimental Design). Focusing on the 295 biochemicals found in at least 10 ingredients, we obtained their nutrient distribution across the food samples using the experimentally measured peak areas. In metabolomics, the peak area for each compound, *n*, in each food, *f*, measures only the ionized concentration, 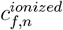. It is unknown what fraction of a compound’s total concentration becomes ionized, preventing peak area from directly measuring the total concentration, *c*_*f*,*n*_. The primary contributors to this loss in efficiency (*E*_*f*,*n*_) are sample preparation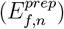, extraction protocols 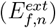, and ionizability 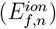 such that 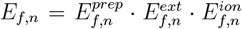 Since our experiments used the same experimental protocol for each food item, *c*_*f*,*n*_ differs from 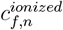 by *E*_*f*,*n*_ where only the food sample matrix provides the largest deviations between measured peak areas across the foods for each compound.

To apply the universal scaling of nutrient concentration to the measured peak areas in untargeted metabolomics, we need 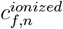 to follow the universal features of nutrient concentration^7^: (i) constant standard deviation (⟨*s*_*n*_⟩), (ii) symmetric distribution, and (iii) translational invariance. A universality rooted in the biochemical mechanisms responsible for the synthesis and consumption of each compound within an organism. If it holds for peak areas, this would mean that we can use the universality to find the total concentrations from peak areas; however, *E*_*f*,*n*_ can break the universality if it greatly varies between food. Yet, we find that (i)-(iii) apply to the peak areas as well, with ⟨*s*_*n*_⟩ = 1.41 ± 0.50. This suggests that

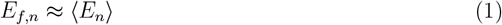

for each compound, where ⟨*E*_*n*_⟩ is the mean efficiency across all foods, conserving the universality. We also find that the peak areas follow log-normal distributions similar to the nutrient concentrations in [7] (SI Universal Scaling Law of Metabolomics). Lastly, we compare the universal scaling observed for nutrients reported by the USDA (Fig 1a) with the biochemicals obtained by our experiments (Fig 1b). We find that the linear standard deviation of peak areas for a biochemical in MassSpec 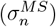 relates to the mean peak area for a biochemical in MassSpec 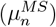 in the log-space via 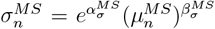, where *α* and *β* are parameters of the power law fit, in excellent agreement with the USDA. These results indicate that the efficiencies of the experiment (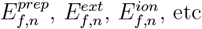etc.) are largely invariant across food sample matrices.

**Figure 1:**
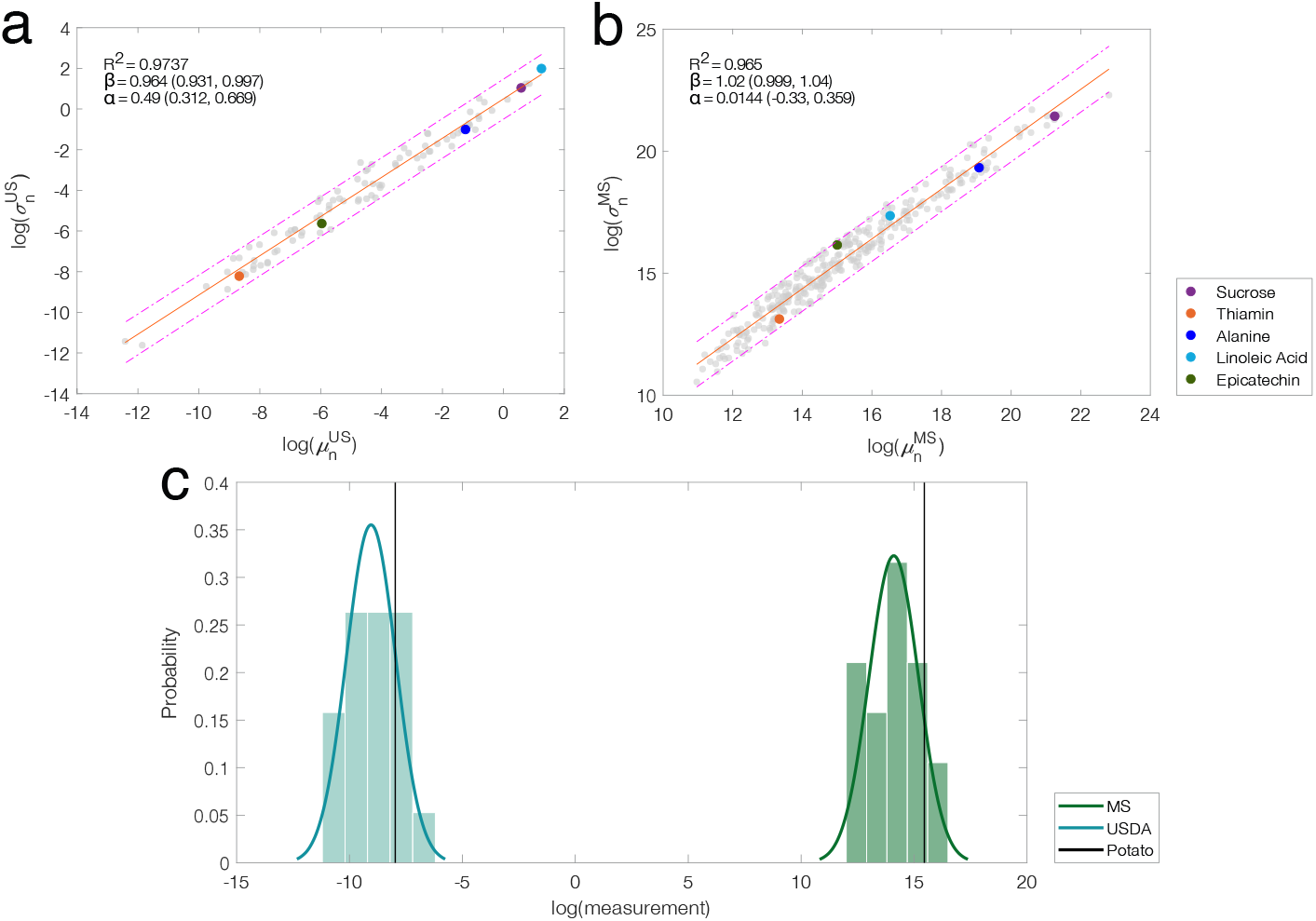
Estimating Concentrations in Untargeted Metabolomics. **(a)** Universal scaling law for nutrients within the USDA subset, relating the standard deviation, 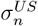 vertical axis, to the average nutrient concentration, 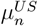 horizontal axis, for 94 nutrients in 510 foods. Dashed lines are confidence intervals. Colored dots are compounds we selected to compare their relative positions between a) and b). **(b)** Universal scaling for our untargeted metabolomics experiments, relating the standard deviation, 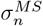, to the mean peak area, 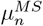, for 295 biochemicals in 20 foods. **(c)** Nutrient distributions using the 19 foods in overlap between the USDA and our experiments for pyridoxine: experiments (peak area, red) and USDA (g/100g, blue). The position of potato is shown in each distribution (black lines).

Next, we test the correlations between the metabolomics variables 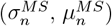 and their respective counterparts of the USDA 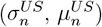. Focusing on 19 foods and 31 biochemicals reported in both datasets (SI USDA – Metabolomics Overlap), we find low correlation (*R*^2^ = 0.566), confirming that ionization efficiency obscures the relationship between the peak areas measured in MassSpec and the concentrations reported in the USDA via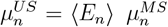. We can, however, estimate the concentration from peak area by leveraging (1) and the universal features (i)-(iii), followed by both metabolomics and USDA, indicating that the position of foods within the nutrient distributions between the datasets is largely conserved. For example, we can see that for pyridoxine (vitamin B6) the relative position of potato is the same in the metabolomics and the USDA distributions (Fig 1c). This suggests that we can use the distance between the individual food measurements and the median in the linear-space to connect the two distributions via

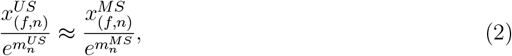

where 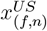 is the concentration of the biochemical for a food from the USDA, 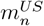 is the mean log concentration of the nutrient in the USDA, 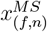 is the peak area of the biochemical in the food from experiments, and 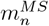 is the mean log peak area of the biochemical in the experiments (SI Proportionality Validation).

To assess the validity of (2) we curate 113 high-quality food-biochemical pairs in overlap between the two datasets by filtering to analytical values with at least 4 measurements (SI Curated Pairs), then estimate 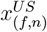 and compare to the reported value in the USDA by calculating prediction error,

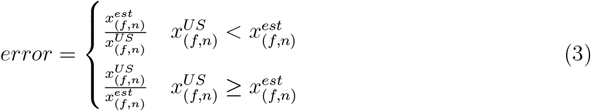

where 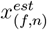 is the estimated concentration as found by Equation (2), 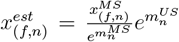 When the estimated and reported concentrations are equal, the prediction error is 1.0 and so values closer to 1.0 is desirable. Using (3), we observe a 3.1 mean error and a 2.4 median error for the 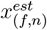 values (Fig 2a, blue line) with 73% of the values below 4.0 error, an outcome comparable with other untargeted metabolomics estimation methods ^8^, which report a 2.0-4.0 mean error. Importantly, this approach should work for any sample where biological and volumetric constraints synergistically determine the concentration (SI Method Guidelines).

**Figure 2:**
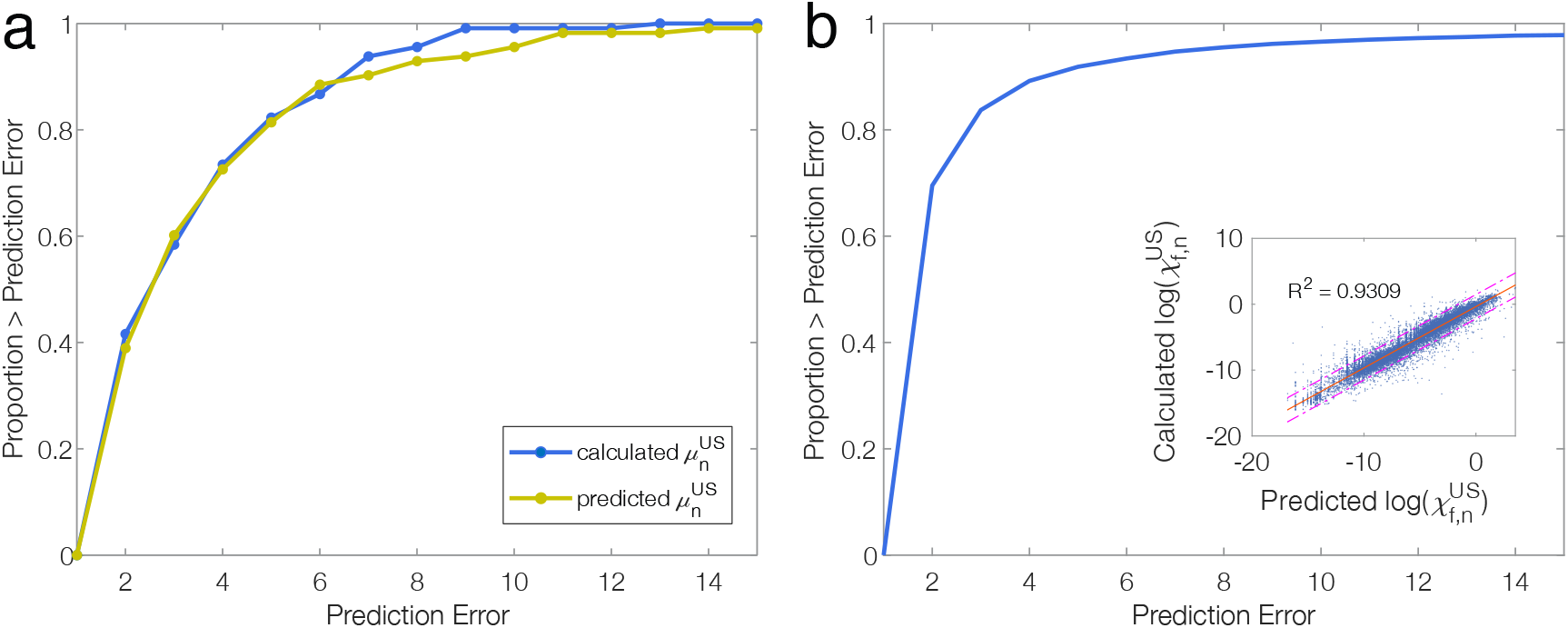
Prediction Error Curves. The proportion of estimated nutrient concentrations in individual food items below specified error values. **(a)** Error curve plots using the curated 113 biochemical-food pairs for the calculated 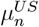 from the USDA (blue) and predicted 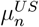 from a XGBoost model (yellow), taking as input the chemical properties of biochemicals (molecular weight, logP, logS, hydrogen bonding inventory, number of charged atoms, non-polar surface area) and phylogenetic lineages of foods (class, order, family, genus classifications). **(b)** Error curve of the Leave-one-out predicted concentrations for the 18,458 biochemical-food pairs in the training data. Inset: the relationship between calculated 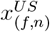 from the USDA and predicted 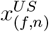 from the XGBoost model. This shows that we can predict the concentrations reported in the USDA with a correlation of *R*^2^ = 0.931 between the predicted and real concentrations.

The methodology (2) requires the concentration 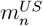 of the biochemicals, known only for nutrients reported in the USDA. To overcome this limitation, we rely on the finding that compound concentrations in bacterial and human cells are correlated with their chemical properties of the compounds^9^, allowing us to infer 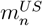 for nutrients not present in the USDA. We created a XGBoost model to predict 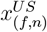, taking as input the chemical properties of biochemicals (molecular weight, logP, logS, hydrogen bonding inventory, number of charged atoms, non-polar surface area) and phylogenetic lineages of foods (class, order, family, genus classifications). Leave-one-out validation of the trained model shows 70% of the 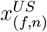 values within 2.0 error and 90% within 4.0 error of the true value (*R*^2^ = 0.931, Fig 2b), confirming the model is accurate (SI Gradient Boosting Methodology).

Using XGBoost to estimate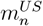, (2) can estimate 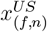 in individual foods using the peak areas, observing a 3.4 mean error (Fig 2a, red line) and 73% of the values below a 4.0 error. XGBoost allows us to determine the concentration of biochemicals not reported in the USDA, but detected in our experiments. For example, while S-allylcysteine in garlic is not reported by USDA, our untargeted experiments allow us to estimate its concentration as 0.158 g/100g. Using FoodMine^10^, we found six published measurements for S-allylcysteine in garlic, with an average at 0.115 g/100g, giving a 1.4 error, demonstrating the possibility of using ML-models to extend the estimated concentrations from (2) beyond the USDA. While promising, the accuracy and generalizability of such models require further study (SI Beyond the USDA).

The proposed methodology offers actionable concentration estimates to complement the standard presence/absence information delivered by untargeted metabolomics, helping managing costs and resources of future studies. We find that our method estimates concentration in untargeted metabolomics with a 3.1 mean error, relying on the universal features of nutrient distributions and offers comparable performance to the current structural similarity method (4.3 mean error) without requiring chemical standards and ionization efficiency prediction method (2.1 mean error) without needing rarely measured ionization efficiencies^8^. Here, we rely on publicly available training data, facilitating the seamless integration of our methodology in the decision-making process of health risk assessments, as seen with established methods^11^ considering food-borne compounds.

## Acknowledgements

We thank Dr. Roger Giese and Dr. Pushkar M. Kulkarni at the Department of Pharmaceutical Sciences, Northeastern University for preparing the food samples. Experiments were performed by Metabolon Inc., led by Nino Esile, Dr. Brian Ingram, and Nathan Testa. This work was partially supported by NIH grant 1P01HL132825, American Heart Association grant 151708, ERC grant 810115-DYNASET, and Rockefeller Foundation grant 2019 FOD 026.

## Data and Code Availability

The data and codes used to develop the methodology are openly available at our GitHub page at https://github.com/Barabasi-Lab/Quantifying-Untargeted-Metabolomics. The raw metabolomics data for the study is available on Metabolights at https://www.ebi.ac.uk/metabolights/MTBLS3319.

## Author Contributions

A.L.B. and G.M. conceived the research. M.S. led the metabolomics experiments and performed the data analysis. A.L.B. and G.M. advised and guided the research. M.S. wrote the manuscript. A.L.B. and G.M. reviewed and edited the manuscript.

## Competing interests

A.L.B. is a scientific founder of Scipher Medicine, Inc., which applies network medicine strategies to personalized drug selection, and Naring, Inc., which applies data science to food and health. All other authors declare no competing interests.

## Notes

https://github.com/Barabasi-Lab/Quantifying-Untargeted-Metabolomics

### Summary of Updates

The manuscript has been updated with material requested by reviewers from a peer review journal.

