## Supplementary Information for "Estimating Biochemical Concentration in Food Using Untargeted Metabolomics"

**Proportionality Validation** **14**

**Method Guidelines** **23**

**Gradient Boosting Methodology** **25**

**Beyond the USDA** **27**

**Model Usage Protocol** **29**

**References** **31**

### Experimental Design

#### Sample Preparation

A selection of 20 produce items were purchased from two local grocery stores (Whole Foods Market and Stop & Shop): apple, banana, basil, black bean, carrot, chickpea, corn, garlic, lettuce, olive, onion, peach, pear, pepper, potato, spinach, soybean, strawberry, and tomato. The items were selected based on genetic and chemical composition data availability, phylogenetic relationships, minimum level of processing, and their consumption patterns within the United States. This selection aims to capture a representative diversity of plant composition in our foods by selecting 13 different phylogenetic families and 19 genus commonly found in the food supply and also accounts for variation in the edible part of plant by selecting fruit (8), leaves (3), roots (3), legumes (3), bulbs (2), and grain (1) foods. With these items, we aim for well-defined nutrient distributions to assess the universal behavior of these distributions[13] in the metabolomics data set. For each produce item, six units were purchased, i.e. six individual apples, six boxes of strawberries, six hands of bananas, six bags of chickpeas. After purchase, the items were prepared for metabolomics analysis[1] in a humidity controlled room while minimizing exposure to light and oxygen to limit biological processes such as browning. The items were prepared through a process of rinsing in water, removing inedible portions (i.e. apple cores, banana peels), and chopping edible portions into 1 $cm^3$ or smaller pieces. All the pieces of the same item were freeze dried at -80 °C for 24 hrs (Catalog No. 10-269-56B from LabConco/Fisher) and then pulverized to yield a fine homogenized powder using a coffee grinder (Kitchen Aid, 170W, Model No. BCG111OBO). The resulting powder is a representative sample over all six units for a specific produce item. The powders were stored at -80 °C to halt the progression of biological processes. For each food, vials were filled with about 200 mg of powder and sent to Metabolon for metabolomics analysis[2]. Argon gas in the vials and dry ice in the shipping box serve to prevent biological processes during shipping.

#### Metabolomics Analysis

Metabolon performed biphasic extraction on a partition of the powder using standard methanol-water extraction. The organic layer was divided into three aliquots, two for positive ionization on a reverse phase platform (UHPLC-CSH C18-HRMS-Orbitrap) at two different parameters and one for negative ionization on the reverse phase platform. The first positive ionization run used acidic conditions, optimized for hydrophilic compounds, with a gradient of a water phase and methanol, 0.05% perfluoropentanoic acid, and 0.1% formic acid phase. The second positive ionization run used acidic conditions, but optimized for hydrophobic compounds, with a gradient of a methanol phase and a acetonitrile, water, 0.05% perfluoropentanoic acid, and 0.1% formic acid phase. The negative ionization run used basic conditions with a gradient of a methanol phase and a water phase both with 6.5 mM ammonium bicarbonate. The hydrophilic layer of the biphasic extraction was used on a HILIC platform (UHPLC-BEH Amide-HRMS-Orbitrap) in negative ionization mode using a gradient consisting of a water phase and a acetonitrile phase both with 10 mM ammonium formate. The MS analysis operated at 35,000 mass resolution and alternated between MS and data-dependent $MS^2$ scans using dynamic exclusion. The scan range was 70-1000 m/z. All 20 produce samples were analyzed on the two platforms with three technical replicates. Metabolon analyzed the resulting spectra using their own internal methodology consisting of four major components, data extraction and peak identification software, data processing tools for QC and compound identification, collection of information and interpretation, and lastly visualizations tools, which generate a chemical composition for each produce item. The raw data was extracted, peak-identified, and quality control processed using a Metabolon designed web-service built platform utilizing Microsoft's .NET technologies. The compounds were identified by comparison to a proprietary library of more than 3,300 purified standards, comparing sample spectra to the reference spectra of these purified compounds. Biochemical identification is further supported by assessing the retention time, mass-charge ratio ($\pm$ 10 ppm), and chromatographic data between the sample and the authentic standards. Quality control was designed to ensure accurate and consistent identification by checking for system artifacts, mis-assignments, and background noise issues. Data analysis was performed to confirm the consistency of peak identification among the various samples. Peaks were quantified using area-under-the-curve. The data was normalized to correct for variation resulting from instrument tuning differences using block correction.

### Data Curation

#### Creating a Plant Only USDA

The Standard Reference and Foundation Foods datasets maintained by the USDA are used as our true concentration values for each food-nutrient pair[3,4]. Since our experimental dataset solely used minimally processed plant-based food items, we filtered the USDA to plant items following the same specifications using keyword search over the food names. Removing foods with animal related keywords such as beef, lamb, fish and with company keywords such as McDonalds, Campbell, Kelloggs. Food names with processing related keywords such as raw, fresh, frozen are kept to ensure only minimally processed items are collected. Combination foods are removed, such as "peas and corn, raw" in order to ensure each concentration relates to a single food. Lastly, the remaining food names are searched through the NCBI taxonomy[5] to find their phylogenetic lineages, if an item was found to not be part of plant kingdom (Viridiplantae), then they were removed. The final list contains concentration information for 510 plant items and 94 compounds.

#### Chemical Properties

For each compound in the USDA and metabolomics experiments, we collected information regarding their chemical properties largely following a previously reported procedure[6]. From PubChem[7], the molecular weight (MW), molecular formula (MF), SMILES, number of hydrogen bond donors (HBD), number of hydrogen bond acceptors (HBA), and number of rotatable bonds (NRB) were collected. Using the MF, we determined the number of carbon atoms (C), number of oxygen atoms (O), number of nitrogen atoms (N), and number of hydrogen atoms (H). The hydrogen bond inventory (HBI) was found by summation of HBD and HBA. The SMILES were used as input into Open Babel[8,9] to determine the number of charged atoms (NCA) at a pH of 7 to simulate a biological environment. The 3D polar surface area (PSA) and 3D nonpolar surface area (NPSA) were calculated using SMILES as input into the rdkit Solvent Accessible Surface Area package[10]. Lastly, the solubility (logS) and partition coefficient (logP) were calculated by ALogPS[11] with SMILES as input.

We performed a correlation of the above properties to the concentrations in the USDA to select the most significant properties. We found that the correlation of the properties improved by roughly 0.1 when adjusting the properties for size of the compound when possible. For example, adjusted HBI is found by HBI/H. NPSA% is found by NPSA/(total surface area). NCA% is found by NCA/(N+O). NRB% is found by NRB/(total number of bonds). The correlation of the chemical properties to concentration data was first used to filter the properties, only using those where the correlation was $\pm 0.4$ or greater. We performed multiple linear regressions, following a previous publication6, with many different combinations of initial chemical properties, reducing down to find the most significant features ($p<0.05$). In addition, the linear regressions were fit to many different subsets of the concentration data (i.e. fitting to primary metabolites versus secondary metabolites or fitting to specific classes of compounds such as amino acids, lipids, and carbohydrates). The chemical properties contributing to the best predictive linear regression models were selected, yielding MW, C, logP, logS, HBI%, NCA%, and NPSA% as our final chemical properties variables to describe each compound in our datasets.

#### Phylogenetic Tree Properties

For each food in the USDA and experiments, a combination of NCBI taxonomy[5] searching and google searching was used to find the scientific names associated with each food item.

Then the NCBI taxonomy was used to collect the full phylogenetic description of the foods, from superkingdom to species. If a food item was not part of the plant kingdom (Viridiplantae), they were removed. Since all foods are in the same kingdom, we only considered phylum and lower. The generic nature of common names in the USDA makes determining subspecies not feasible and often the species could not be determined, therefore, we collected from phylum to genus for all foods. There is heterogeneity the tree descriptions for each food, some containing subfamilies and some containing a multitude of clades. We only selected levels where the 90% or more of the foods had a description. This left us with the most common phylogenetic tree categories: phylum, class, order, family, and genus collected for all food items.

#### Phenol-Explorer

We obtained Phenol-Explorer[12] to improve our resolution of the secondary metabolism and increase the accuracy of our models. The dataset went through the same process as described in Section Creating a Plant Only USDA. The chemical and phylogenetic properties for the biochemical-food pairs are collected following the descriptions in Sections Chemical Properties and Phylogenetic Tree Properties respectively. The remaining Phenol-Explorer dataset contains 230 compounds, adding resolution to the secondary metabolites for the foods within the USDA. 878 food-nutrient pairs are found and added to the USDA when training the gradient boosting algorithm.

#### USDA-Metabolomics Overlap

The average concentration of each food-nutrient pair is obtained from the Standard Reference and Foundation Foods[3,4]. The overlap of the USDA datasets to the experimental metabolomics results is determined by matching both food and nutrient information. First, the datasets are matched by the foods, using food descriptions within the USDA and the sample preparation used in the experiments; for example, apple samples are prepared with skin and thus are matched to “apple, raw, with skin” within the USDA. Only the compounds detected in the experiments and contained in the USDA are included, this is achieved by using PubChem[7] to find InChIKeys to match the compounds between the datasets. The overlap creates a curated dataset that has 19 food items and 49 compounds. Lastly, we remove compounds present in 10 or fewer food items, creating a finalized list of 31 compounds. For the USDA, we obtained the reported average concentration for each nutrient in each food item. For the experiments, we have the average peak area over three technical replicates for each biochemical in each food item.

#### Curating Quality Pairs

The quality of the data in the USDA is highly variable in the methods used to quantify compounds as well as the number of data points for a compound within a food, so in order to assess the accuracy of the method at determining concentrations; we want to reduce the variability originating from the USDA. Initially we have 19 foods and 31 compounds, making a total of 589 biochemical-food combinations; however, not all compounds are present in all foods of the USDA or in the peak areas of metabolomics. If the USDA has 0 concentration or if metabolomics has 0 peak area, then the biochemical-food pair is removed. The resulting list has 391 biochemical-food pairs remaining. The remaining pairs have a 11.1 mean error. The large error comes from a combination of method used and number of data points in the USDA. If calculating the mean error for specific methods, we can see how quality is affected. For example, pairs labeled with “Based on another form of the food or similar food” yield a 25.0 mean error while pairs labeled with “Analytical or derived from analytical” yield a 7.6 mean error. Next, we considered the number of data points impact by setting a threshold, only including those at or above the specified data point threshold. If the threshold is 1 data point the mean error is 6.3, set threshold to 4 data points the mean error is 5.5, and at 8 data points the mean error is 4.9. The more measurements made for a pair, the more accurate our estimation. For this paper, in order to remove lower quality data and provide a better assessment of our estimations, we only used data marked as analytical (source codes 1, 6, 12, and 13) and food-nutrient pairs which had 4 or more data points. Once controlling for method type and number of data points, we had 113 pairs. Due to the filtering, the finalized list does not represent each food or nutrient equally, so some are represented more than others. In Table S1, the breakdown of the food representation is shown. In Table S2, the breakdown of the compound representation is shown.

Table S1: **Food in Filtered USDA.** The number food-nutrient pairs attributed to each food item in the filtered USDA of 113 pairs used to assess the accuracy of our estimated concentrations.

| **Food** | **Count** |
|---|---|
| apple | 6 |
| banana | 5 |
| basil | 1 |
| carrot | 7 |
| chickpeas | 2 |
| corn | 18 |
| lettuce | 7 |
| onion | 6 |
| peach | 7 |
| pear | 5 |
| pepper | 6 |
| potato | 6 |
| soybean | 2 |
| spinach | 24 |
| strawberry | 6 |
| tomato | 6 |

Table S2: **Nutrients in Filtered USDA.** The number food-nutrient pairs attributed to each compound in the filtered USDA of 113 pairs used to assess the accuracy of our estimated concentrations.

| Nutrient | Count |
|---|---|
| alanine | 2 |
| alpha-tocopherol | 9 |
| arginine | 2 |
| aspartic acid | 2 |
| fructose | 12 |
| glucose | 12 |
| glutamic acid | 2 |
| glycine | 2 |
| histidine | 2 |
| isoleucine | 2 |
| leucine | 2 |
| lysine | 2 |
| methionine | 2 |
| nicotinic acid | 11 |
| phenylalanine | 2 |
| proline | 2 |
| pyridoxine | 14 |
| serine | 2 |
| sucrose | 11 |
| thiamin | 10 |
| threonine | 2 |
| tryptophan | 2 |
| tyrosine | 2 |
| valine | 2 |

### Universal Scaling

#### Universal Scaling of USDA

Using the subset of the USDA described in Data Creation, Creating a Plant only USDA subsection as input, the universal features of each nutrient distribution across the foods in the log-space were detected: (i) constant standard deviation ($\langle s_n \rangle$), (ii) symmetric distribution, and (iii) translational invariance[13]. Here we confirm that the universal features (i)-(iii) hold with $\langle s_n \rangle = 1.64 \pm 0.54$ for the smaller subset of the USDA data, representing 12% of the foods from the initial study. Again, we observe the same dependence of the linear standard deviation ($\sigma_n^{US}$) on the mean nutrient concentration ($\mu_n^{US}$) in the log-space, captured by the power law fit $\sigma_n^{US} = e^{\alpha_\sigma^{US}} (\mu_n^{US})^{\beta_\sigma^{US}}$ ($R^2 = 0.974$, Main text Figure 1a).

#### Universal Scaling of Metabolomics

Here we provide a more general assessment supporting that metabolomics data follows the universal scaling in [13], similarly to the USDA concentration data. We obtained over 600+ identified compounds from the experiments over the 20 food items tested, the list contains labeled compounds (those with known names and structures) and unlabeled compounds (those unnamed and without a structure). We observe that the peak area probability distributions across foods, Q($m_{peaks}$), obtained from metabolomics follow similar patterns as those previously reported for the USDA[13] (Fig S1a). The distributions are largely symmetric in the log-space with similar shapes that do not depend on the average peak area of the compound. There is deviation in shape between compounds as a result of the small number of foods within the metabolomics dataset. Furthermore, we see that the logarithmic standard deviation ($s_{peaks}$) of the MassSpec peaks is independent of the logarithmic mean ($m_{peaks}$), the same observation is found within the USDA data (Fig S1b). In fact, the logarithmic standard deviation distributions of the MassSpec peaks ($s_{peaks}$) and USDA nutrients ($s_{nutrients}$) have large overlap, showing that the mean $s_{peaks}$ is about equal to the mean $s_{nutrients}$ of the two datasets. The universal scaling can be obtained using all the compounds provided as shown in Fig S1d, yielding $R^2 = 0.962$ for the dependence of $\sigma_{peaks}$ on $\mu_{peaks}$. Both known and unknown compounds follow the universal scaling.

Next, we evaluated the performance of four probability distributions: lognormal, gamma, Weibull, and Gaussian. The best fit for each distribution to the peak areas in the log-space over all 20 food items for 625 compounds is found in order to derive the scale and shape parameters for each distribution and the location parameter for the Gaussian. This includes all compounds identified, both known and unknowns, not just the compounds in overlap with the USDA. Model data is generated for each probability distribution by random sampling from the best fit and then compared to the real data using the Kolmogorov-Smirnov (K-S) test. We find that the cumulative probability distributions of the K-S distance over the compounds as shown in Fig S1e. We see that the lognormal distribution provides the best fit slightly over Weibull distribution and Gamma distribution while Gaussian is the worst. This is consistent with our previous findings on the USDA data that lognormal provides the best fit[13]. However, due to the small number of foods within the metabolomics data, the difference of lognormal to the other distributions is less prominent.

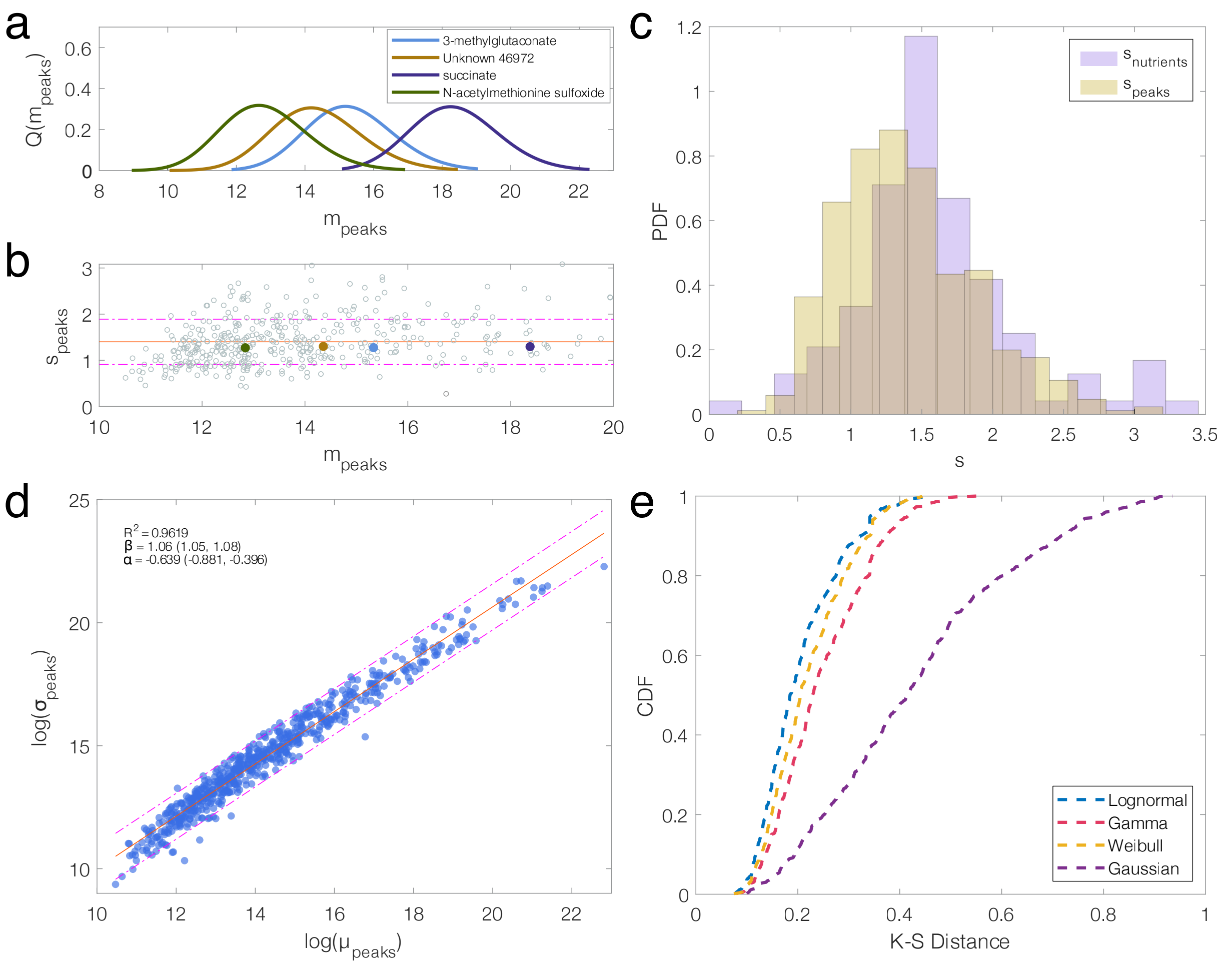

Figure S1: **Metabolomics Universal Scaling (a)** The peak area probability distribution, Q($m_{peaks}$), for five compounds over the 20 foods in the experiments. The distributions are similar though subject to deviation due to the small food panel. **(b)** Logarithmic standard deviation of the peaks ($s_{peaks}$) to the logarithmic mean ($m_{peaks}$). Colored points represent the position of the compounds in a). **(c)** Histograms of the probability density function for the USDA ($s_{nutrients}$) and MassSpec ($s_{peaks}$). **(d)** The dependence of $\sigma_{peaks}$ on $\mu_{peaks}$ for all the compounds detected within the experiment, containing both compounds with known structures and those that are unknown. **(e)** Fit performance of four probability distributions using Kolmogorov-Smirnov distances for assessment.

### Proportionality Validation

From the main text, we see that the universal features (i)-(iii) allow the position of the individual food items within the nutrient distributions to be largely conserved between the USDA and metabolomics experiments. This allows us to establish a proportionality based on the distance to the mean in the log-space (as seen in Figure 1c of the main text) in order to circumvent the effects of ionization efficiency,

$$log(x_{f,n}^{US}) - m_n^{US} \approx log(x_{f,n}^{MS}) - m_n^{MS} \tag{1}$$

where $m_n^{US}$ is the mean log nutrient concentration in the USDA, $m_n^{MS}$ is the mean log peak area in the experiments, $x_{f,n}^{US}$ is the concentration for a single compound in a single food item in the USDA, and $x_{f,n}^{MS}$ is the peak area for a single compound in a single food item in the experiments. From (1), we can substitute both $x_{f,n}^{US}$ and $x_{f,n}^{MS}$:

$$(m_n^{US} + Z_{f,n}^{US} s_n^{US}) - m_n^{US} \approx (m_n^{MS} + Z_{f,n}^{MS} s_n^{MS}) - m_n^{MS} \tag{2}$$

where $s_n^{US}$ is the standard deviation in the log-space for the USDA, $s_n^{MS}$ is the standard deviation in the log-space for the experiments, $Z_{f,n}^{US}$ is the Z-score of the food-nutrient pair in the USDA, and $Z_{f,n}^{MS}$ is the Z-score of the biochemical-food pair in the experiments. This simplifies to:

$$e^{Z_{f,n}^{US} s_n^{US}} \approx e^{Z_{f,n}^{MS} s_n^{MS}} \tag{3}$$

From (3) we find the Z-ratio, $e^{Z_{f,n}^{US} s_n^{US}} / e^{Z_{f,n}^{MS} s_n^{MS}} \approx 1$, to assess the approximation. While (3) can be further simplified, Z-scores can be negative and it is important to remove the effects of sign, especially for foods near the mean. In Figure S2, we see that the Z-ratios are largely near 1, in agreement with the approximation found in (3), with a median of 1.8, further supporting the proportionality between the distributions in (1). The proportionality in the log-space is translated to the linear space by taking the exponential of (1) resulting in:

$$\frac{x^{US}_{(f,n)}}{e^{m^{US}_n}} \approx \frac{x^{MS}_{(f,n)}}{e^{m^{MS}_n}}, \tag{4}$$

as seen in the main text. Due to the lognormal distribution fit of the data, Equation (4) translates the log-space into the linear space via a property of lognormal distributions, where the median of the data in the linear space is equal to the exponential of the mean in the log-space, i.e. $\mathrm{Med}(x^{US}_n) = e^{m^{US}_n}$ and $\mathrm{Med}(x^{MS}_n) = e^{m^{MS}_n}$. Here, $x^{US}_n$ is the concentrations for the compound in all foods and $x^{MS}_n$ is the peak areas for the compound in all foods.

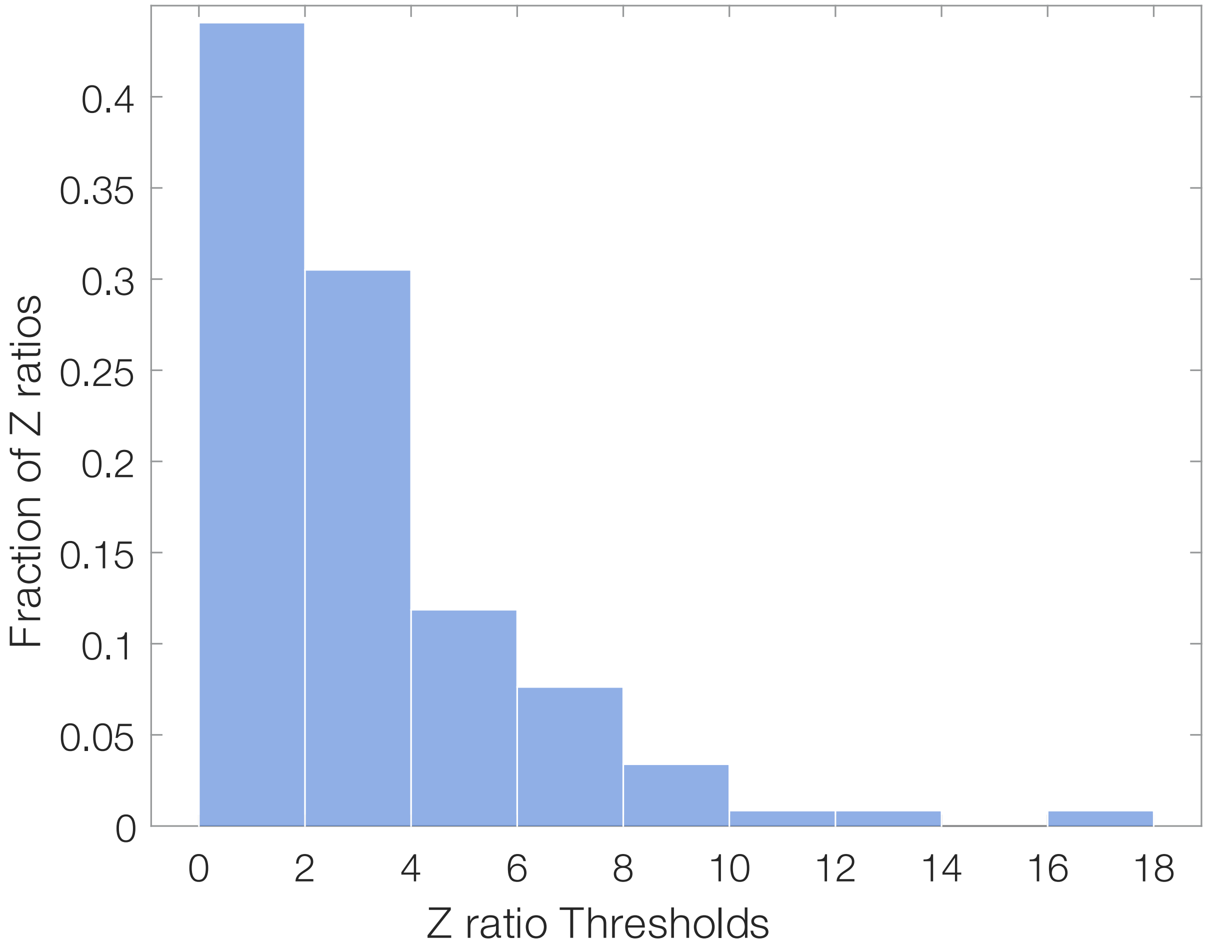

Figure S2: **Z-ratio approximation**. A histogram of the Z-ratio of the curated overlap compounds and foods. The approximation holds if Z-ratio is near 1, we find that the median is 1.8 for the dataset.

#### Using the Proportionality for Estimation

As seen in the previous section, there exists a proportionality, Equation (4), between the USDA values and the metabolomics values which can be leveraged to estimate the concentration of a compound within each food item. The accuracy of Equation (4) is heavily dependent on the quality of the data used within the analysis. For example, as observed in Supplementary Information section Curating Quality Pairs, we observed that the number of experimentally measured data points within the USDA for a biochemical-food pair can impact the accuracy through $x^{US}_{(f,n)}$. Each value within Equation (4) needs to be carefully established to ensure sufficiently low prediction error. The number of foods within the USDA will impact the $m^{US}_n$ value, so creating large subset of the USDA to determine this value will be needed. In Supplementary Information section Creating a Plant Only USDA, we can see that a large data set can be made to ensure an accurate value. Indeed, these same considerations are essential for the metabolomics side of Equation (4) as well. Here, we define $\text{Mratio} = x^{MS}_{(f,n)}/e^{m^{MS}_n}$, the right-hand side of the equation, and look into the impacts of the number of foods on its value. In order to make an assessment on the number of foods required to provide accurate estimation, we take a look at the biochemicals in pear and observe how their Mratios change with various sizes in the number of foods (Fig S3). This is achieved by randomly selecting between the 19 others foods within the experiments and adding pear. For example, to find the Mratio using three foods, we randomly select two foods that are not pear then add pear as the third food. This ensures that pear is present in all Mratio calculations. This is repeated to find the Mratio for 140 random sets containing 3 of our foods. Then this is completed for sets of 3, 5, 7, and so on of our foods. We observe in Figure S3a that for compounds in pear, the Mratio exhibits instability at a lower number of foods and becomes more stable with the addition of foods. The change in the observed Mratio decreases as the number of total foods used increases. The shift of Mratio with increasing number of food is dependent on the peak area of the compound in pear. For example, epicatechin in pear is large, on the right side of the nutrient distribution similar to other Rosaceae foods; however, a small number of foods yielded the possibility that other Rosaceae food items (i.e. apple, peach, strawberry) are present in the random set, and thus making the amount of epicatechin look more normal. As the number of foods increases, the exceptional amount of the compound in pear becomes more apparent, increasing the Mratio value. This same effect is occurring for tryptophan, yet in the opposite direction where the addition of more foods reveals that the amount of tryptophan in pear is smaller than most foods. This uncertainty in the Mratio is clearly observed in the standard deviation per compound in pear (Fig S3b). We can see that at three food sets the standard deviation is large, reaching about a 6% swing for glucose due to the smaller set not representing food diversity well. With the use of 11 food items, the standard deviation of Mratio has halved and by 17 food items all five compounds plotted are nearing a 1% deviation. With this in mind, these data suggest that around 11-13 food items the Mratio tends to reduce its variability and start to stabilize in value. The food items selected also need to be varied, covering a wide range of edible phylogenetic families and genus in order for Mratio to accurately represent the food. In addition, the part of plant should also be considered when making the selection as the part of plant impacts the compositional makeup of a food in addition to the phylogenetic lineages. To match the quality of this study, likely 19 phylogenetic genus spanning fruit, leaves, root, and legume will be necessary.

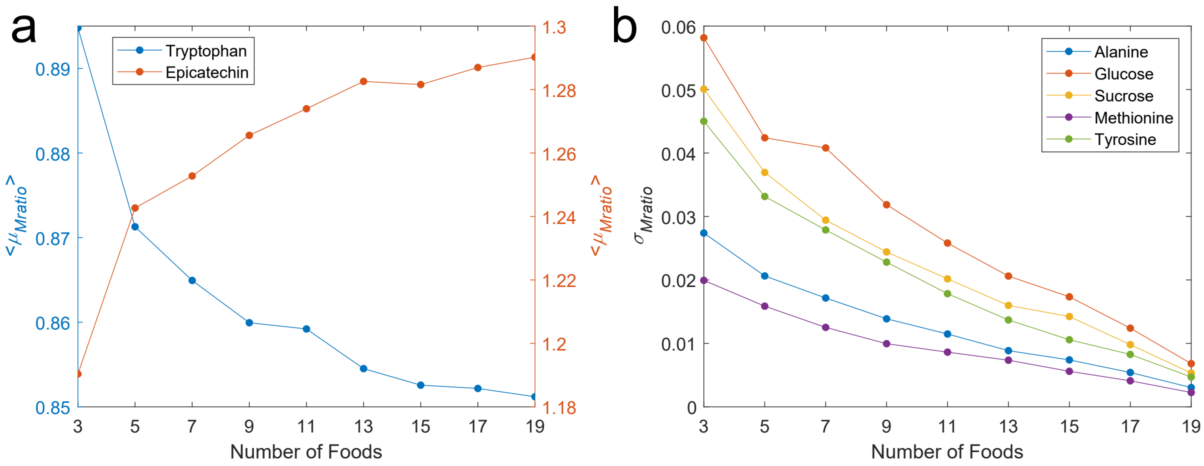

Figure S3: **The Metabolomics Ratio Assessment**. A random number of foods is selected in addition to pear to study the effect of the number of foods used to calculate the metabolomics ratio of compounds in pear (number of random sets = 140 for each point except when number of foods = 19). **a)** The average metabolomics ratio of tryptophan (blue, left-hand y-axis) and epicatechin (red, right-hand y-axis) within pear. The averages are calculated over the random sets with a specified number of food items. **b)** The standard deviation of the metabolomics ratio over the random sets at each specified number of food items for five compounds detected in pear.

#### Proportionality and Linear Regression

In the main text, we showed that the position of individual food items within the nutrient distributions, due to the universal features (i)-(iii), is largely conserved between the USDA concentrations and metabolomics peak areas. This also suggests that there exists a linear relationship in the log-space between the concentrations and peak areas for each compound across the food items,

$$log(x_n^{US}) = \beta log(x_n^{MS}) + \alpha, \tag{5}$$

where $x_n^{US}$ is a vector of the USDA concentrations for compound $n$ for all foods and $x_n^{MS}$ is a vector of the metabolomics peak areas for compound $n$ for all foods. The proportionality shown in Equation (4) can be rearranged into a linear form:

$$log(x_{f,n}^{US}) = log(x_{f,n}^{MS}) - m_n^{MS} - m_n^{US}, \tag{6}$$

where $x_{f,n}^{US}$ is the reported concentration of compound $n$ in food $f$ within the USDA, $x_{f,n}^{MS}$ is the peak area of compound $n$ in food $f$ within the metabolomics experiments, $m_n^{US}$ is the mean log concentrations from the USDA, and $m_n^{MS}$ is the mean log peak areas from metabolomics. Here, we see that Equation (6) and Equation (5) are functionally similar when $\beta \approx 1$, and suggests that

$$\alpha \approx m_n^{US} - m_n^{MS}. \tag{7}$$

Indeed, this equivalency is further supported by Ordinary Least Squares (OLS) where the intercept, $\alpha$, is the average response, defined as:

$$\alpha = \frac{1}{N}\sum_{f=1}^{N} log(x_{f,n}^{US}) - \beta log(x_{f,n}^{MS}), \tag{8}$$

containing contributions from each food $f$ in the total number of foods studied ($N$). We see that $log(x_{f,n}^{US}) = m_n^{US}$ and $log(x_{f,n}^{MS}) = m_n^{MS}$. Using Equation (5), we calculate the

$\alpha$ for each compound in overlap between the USDA and the experiments. Then, we find the ratio, $\alpha/m_n^{US} - m_n^{MS}$ according to Equation (7). When the ratio is near 1.0, then the approximation holds, showing that the linear regression method and the proportionality method are largely equivalent. As we can see in Figure S4, the ratio is approximately 1.0 for most compounds. The compounds greatly above 1.0, alpha-tocotrienol and beta-cryptoxanthin, are due to small $N$ and thus lack an adequate number of foods necessary for higher accuracy, as discussed in the section Using the Proportionality for Estimation. We can explore the similarities between the two methods in more details, for example with sucrose. There are 12 food items with reported concentrations in the USDA and peak areas from the experiments. In addition, 11 of the measurements in the USDA are determined to be high quality as discussed in section Curating Quality Pairs. We find an $R^2 = 0.81$ for the linear regression in the log-space between the USDA and MassSpec values. In addition, the Spearman correlation is 0.90 between the USDA concentrations and experimental peak areas in the log-space, showing that the rank of the foods between the distributions is conserved. Therefore, the universal features (i)-(iii) of nutrient distribution provide an explanation for why the proportionality method and the linear regression method can yield reasonable estimations of compound concentrations in food items. The universal features are essential to understand when considering their success in estimating concentrations.

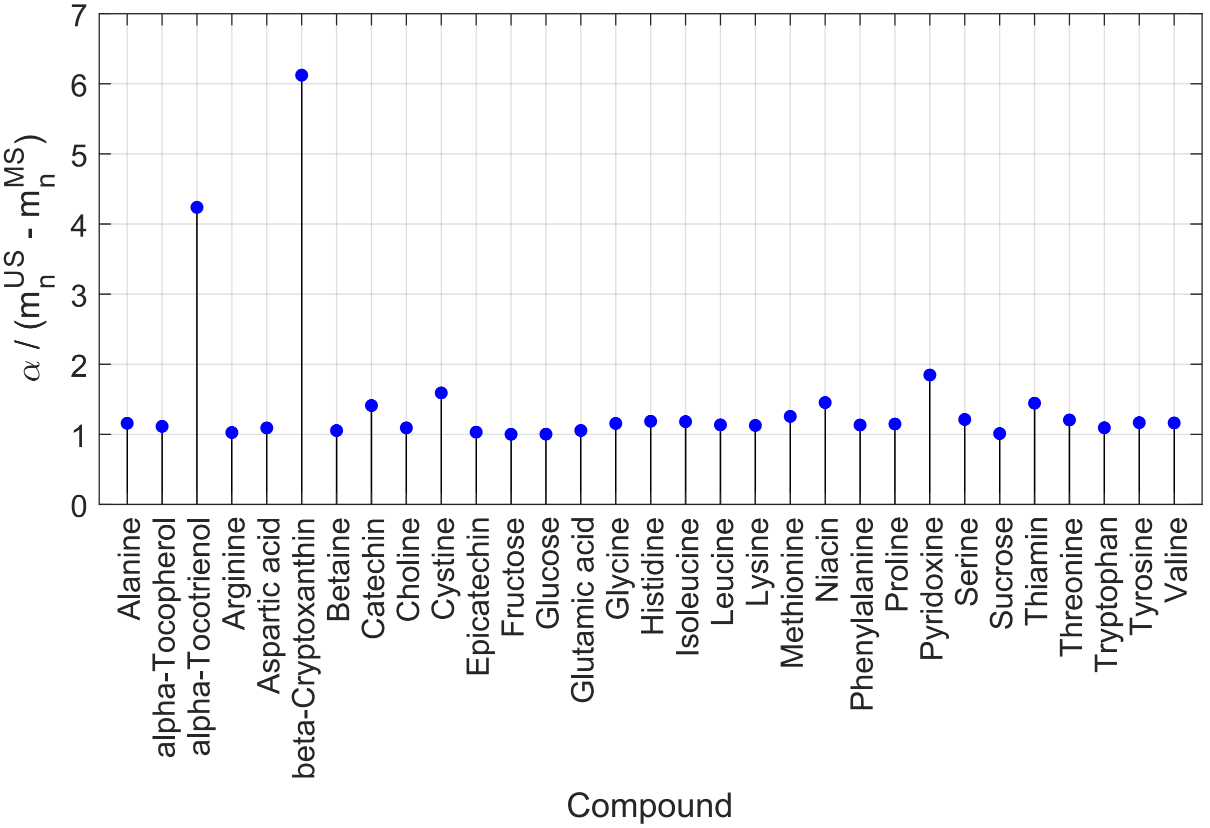

Figure S4: **Proportionality and linear regression**. The ratio of the proportionality found through the universal features ($m_n^{US} - m_n^{MS}$) and the intercept of linear regression in the log-space ($\alpha$). The ratio for each compound is approximately 1.0 in agreement with Equation (7).

### Method Guidelines

Our approach to estimating concentration in untargeted metabolomics is based on the universal scaling of nutrient concentrations[13] which is based on the biological and volumetric constraints imposed on biochemical concentration in the synthesis and consumption of the biochemicals in cellular organisms. These biological processes establish the universal features of nutrient concentration for all biochemicals: (i) constant standard deviation, (ii) symmetric distribution, and (iii) translational invariance. We showed that the universal scaling holds for the whole food supply (USDA Standard Reference), only raw ingredients, different chemical classes such as plant secondary metabolites, and to various other databases beyond the USDA. The method is robust and we expect it to hold for any biological sample where biochemical reactions and volumetric constraints govern concentrations. Indeed, we find the universality on the whole food supply, covering both animal and plant products. Even many processed foods and complex dishes follow universal scaling. The potentially wide applications of universal scaling can make it difficult to distinguish when our method to estimate concentrations in untargeted metabolomics is applicable. To simplify the assessment, researchers should pose two questions regarding the possible application: 1) Are the compounds modulated biochemical reactions within the organism? and 2) Are the compounds bound by volumetric constraints?

#### Processing Alterations

For processed foods, we investigated biochemicals affected by differing process and fortification and observed deviations from the expected log-normal distribution with many outliers altering the shape of the distribution[14]. The addition of fortifications, e.g. adding vitamin D to orange juice, or additives, e.g. adding sodium for preservation, can violate the universality. However, our analysis, based on the logarithmic nature of the nutrient variability, minimizes the impact of these deviations, allowing for us to uncover the dominant patterns of variance. Furthermore, whenever fortified biochemicals or additives are included in food following the concentration of natural compounds by a renormalization factor, log-normality would be found as a consequence. In summary, while the consistent scale of nutrient fluctuations across the food supply is constrained by biochemical principles, the nutrient contents for a given food display different patterns of outliers which are indicative of its degree of processing. This disruption can be used to measure the level of processing for each processed food[15]. Yet, these processing alterations can make estimated concentrations within untargeted metabolomics for processed foods unreliable.

#### Effect of Boundary Constraints

We found that the universal scaling is impacted by the behavior of nutrients with the highest concentrations[13]. Biochemicals with concentrations near the sample mass are found to have their concentrations modulated by the sample mass rather than the universal scaling, their abundance and variability driven by the fixed mass of the sample. We observed this behavior in the whole food supply, USDA Standard Reference, with a sample mass of 100g and measured macronutrients: Carbohydrates, Total Fat, Protein, Total Sugars, Total Monosaturated Fatty Acids, and Total polysaturated Fatty Acids. While the universality holds for the individual micronutrients part of each macronutrient, each under biochemical constraints, their total sum yields a distribution in proximity to the sample mass. This effect is also observed with water, a single biochemical, whose concentration does not find convergence due to its proximity to these boundary constraints.

#### Example of Violations to the Methodology

If a datasets is determined to have biochemical and volumetric constraints, it is advised that researchers perform an analysis for the features of the universality (i-iii) as discussed in Section SI Universal Scaling of Metabolomics.

Here we will use a dataset of pollutant concentration in groundwater[16] to demonstrate a case where the universal features are violated and thus our method cannot be used to estimate concentration in untargeted metabolomics. The dataset contains 31 groundwater samples from 9 springs and 22 abstraction wells collected by the Swiss National Groundwater Monitoring NAQUA, operated by the Federal Office for the Environment. The groundwater is collected in samples of 1000mL and the concentrations of 29 pollutants are reported in ng/L. It is clear that biological processes are unlikely regulate their concentrations. Instead, it is more likely driven by governmental regulations, sample location, i.e. proximity to a farm where a particular pesticide is used, and the time of year the sample is collected. Therefore, we can use this dataset to demonstrate how the analysis of universal features can show violations. Since our analysis is statistical in nature, we filter the pollutants to those detected in four or more groundwater samples and only pollutants annotated with an authentic reference standard. The remaining 12 pollutants were detected in $24 \pm 6$ groundwater samples while on average each groundwater sample had $8 \pm 3$ pollutants detected. We see that (ii) is often violated for the pollutants where the concentrations provide discontinuous distributions with near uniform behavior (Figure S5). Interestingly, (iii) is difficult to assess due to all pollutants spanning a similar range in concentration, possibly driven by use regulations. (i) is violated in that the standard deviation in the log-space is not normally distributed around a constant value (Figure S5). Lastly, we generate a KS-distance plot which shows that log-normality is not the best fit for the entire dataset (Figure S5). In summary, we find that the pollutants in the groundwater do not follow universal scaling, breaking features (i) and (ii), and that there are other distribution model that fit the dataset better than lognormal.

### Gradient Boosting Methodology

The gradient boosting model was created using python packages scikit-learn[17] and scikit-optimize[18]. Principal component analysis (PCA) is performed on the chemical properties (MW, C, logP, logS, HBI%, NCA%, and NPSA%) due to significant collinearity between

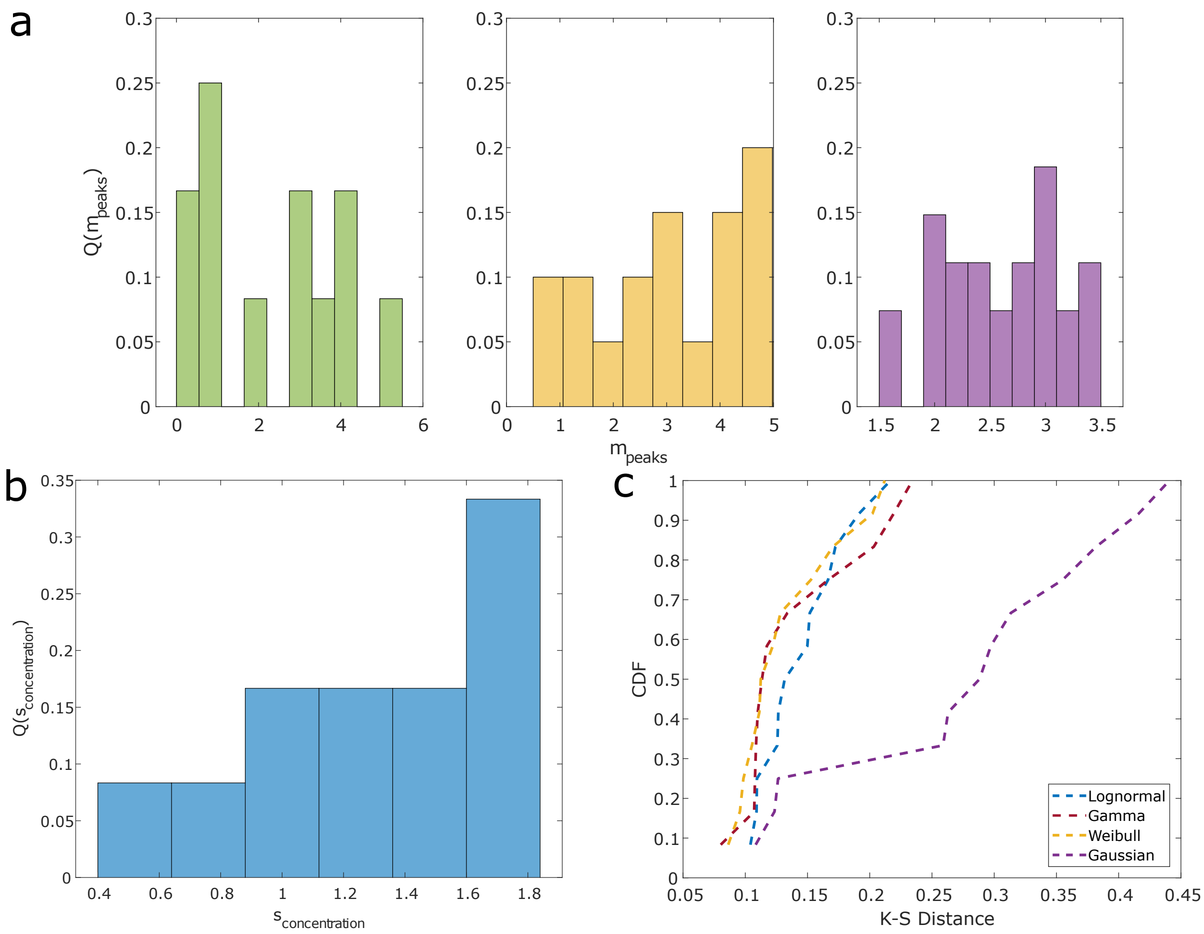

Figure S5: **Assessing Universal Scaling on Groundwater (a)** The concentration probability distribution of pollutants, Q($m_{concentration}$), for 3 compounds over the 31 groundwater samples, Chlorothalonil-TP SYN507900 (left), Metolachlor-TP NOA413173 (middle), and Terbuthylazine-TP CSCD648241 (right). The distributions have different shapes for each pollutant. **(b)** The probability distribution of the logarithmic standard deviation of the pollutant concentrations ($s_{concentration}$). The pollutants lack a constant $s_{concentration}$. **(e)** Fit performance of four probability distributions using Kolmogorov-Smirnov distances. Lognormal does not have the best fit for the dataset.

chemical properties. The top 4 components are selected due to covering 96.80% of the explained variance and how feature importance changes as components are removed when training the gradient boosting model. Multiple correspondence analysis (MCA) is performed on the phylogenetic tree properties (class, order, family, and genus) in order to reduce the number of dimensions for processing speed. We took the top 13 components covering 14.35% of the explained variance. Since experiments and the USDA can have different compounds and a different number of foods, individual PCAs of the training and test dataset will yield components composed of different feature spaces. In order to project both into the same feature space, the PCA should be performed over all compounds and foods in both datasets. Scikit-opt optimized the hyperparameters through a Bayesian cross validation search through a search space controlling max_depth (3-10), max_leaf_nodes (2-50), learning_rate (0.01-1), number_estimators (10-200), and K-Fold = 5 with shuffle set to true. It performed 100 iterations over the search space to select the optimal hyperparameters. For the USDA only model the final hyperparameters are max_depth = 10, max_leaf_nodes = 43, learning_rate = 0.1235, and number_estimators = 95. For the USDA and Phenol-Explorer model the hyperparameters are max_depth = 10, max_leaf_nodes = 50, learning_rate = 0.09724, and number_estimators = 200. The $R^2$ is tracked by scikit-opt for each K-Fold and shows that $R^2 = 0.898 \pm 0.002$ over the 5-Folds for the USDA model and $R^2 = 0.885 \pm 0.004$ over the 5-Folds for the USDA and Phenol-Explorer model. The models were both validated by leave-one-out methodology and we collected the mean squared error (MSE) for each. For the USDA model MSE is $0.901 \pm 2.85$ (0.170 median). For the USDA and Phenol-Explorer MSE is $1.04 \pm 3.29$ (0.184 median). Further inspection of the MSE distributions reveals that a few trials are skewing the mean with 80% of the MSE values below the respective means in both models. We use the USDA and Phenol-Explorer model in the paper since it provides a better prediction error curve than the USDA model alone. The USDA contains data points for 17,753 biochemical-food pairs, dominated by primary metabolites, while the Phenol-Explorer contains data points for 878 biochemical-food pairs, dominated by the secondary metabolites. Therefore, the training data set contains 18,458 total biochemical-food pairs. The improved prediction error curve for the combination model is likely because the model is more generalizable, more able to predict both primary and secondary metabolites.

### Beyond the USDA

As shown in Proportionality Validation section, to estimate the concentration of a compound in metabolomics through (4), we require the log-space average concentration from the USDA ($m_n^{US}$). This limits the application of our approach to compounds with sufficient data to determine $m_n^{US}$. However, using the gradient boosting model, we can estimate the concentration of compounds detected by metabolomics that are not reported within the USDA, circumventing the data limitations of the USDA. We designed an approach where the 5-folds of the USDA and Phenol-Explorer model, stratified by compound class, each predict the concentration for a chosen compound in all foods within the dataset. Since the biochemical composition is specific, it is unreasonable to assume the compound is present within all the foods. The model will always predict a concentration since it has no knowledge of the biochemical composition specificities. Therefore, we need to set a minimum threshold to the predicted concentrations, below which the prediction is set to zero. For this demonstration, we set the threshold to be -11 g/100g in the log-space, the first quartile of the minimum concentrations reported for each compound over all foods. If a fold predicted the concentration below the threshold, then its prediction was set to zero. We find the average predicted concentration over all 5-folds to find the predicted concentration for a single food item. If the majority of the folds are set to zero, then we predict food item does not contain the compound. This allows us to predict $m_n^{US}$ by finding the average over all food items with predicted non-zero concentrations and input into (4) to calculate the concentration of compounds in the metabolomics dataset which have no USDA reported data.

Our values from (4) are compared to measured values in the literature found using FoodMine[19]. We compared the compounds in overlap for garlic between the experiments and the literature search. The FoodMine results were filtered to remove entries that used garlic as cooked, processed, or prepared as part of a recipe. All data that is measured from different growing regions were considered to calculate the average concentration in garlic. There are five overlap compounds that meet these criteria: fumaric acid, caffeic acid, ferulic acid, S-methylcysteine, and S-allylcysteine. Using the FoodMine measurements as the true values, we calculate the prediction error for the estimated concentrations. For fumaric acid and caffeic acid, we observe a 12.1 error and 10.6 error respectively. The larger error is due to the number of measurements being below the 4 measurement threshold used for the USDA. Ferulic acid with 5 measurements yielded a 1.6 error. S-methylcysteine and S-allylcysteine both have 6 measurements, and provide a 1.2 and 1.4 error respectively. This result is similar to the USDA data, in that the number of measurements is important to set a sufficiently informed average concentration for comparison with our predictions. With 4 or greater measurements, the accuracy of our predictions improves. This methodology shows that we can use the XGBoost model to predict a compound's expected $m_n^{US}$ and provide estimations of concentration to metabolomics data even when there are no reported measurements in the USDA.

### Model Usage Protocol

From these results, we can design a protocol for use of this model. In order to predict the mean total concentration of a compound within a specific food, researchers should follow SI Sections Chemical Properties and Phylogenetic Tree Properties to obtain the required features and then follow SI section Gradient Boosting Methodology to apply the appropriate transformations for input in the model. Give this to the model for the output prediction of the mean total concentration of that compound in the selected food. Since the model is sparse in terms of compound-food pairs, we recommend researchers to consider potential gaps in the model's training. For example, if the compound is not found within the USDA or Phenol-Explorer or differs greatly in chemical properties/chemical structure from those in the training data, then additions to the training dataset might be necessary to improve results, which will require retraining the model. This will require adding a multiple compound-food pairs with compounds similar to the one of interest. The same applies to the food items since the training databases largely collect on commonly consumed foods, there can be gaps within the phylogenetic tree which might need to be corrected if the food of interest is far genetically from common foods.
